# Thermodynamic consistency of autocatalytic cycles

**DOI:** 10.1101/2024.10.11.617739

**Authors:** Thomas Kosc, Denis Kuperberg, Etienne Rajon, Sylvain Charlat

## Abstract

Autocatalysis is seen as a potential key player in the origin of life, and perhaps more generally in the emergence of Darwinian dynamics. Building on recent formalizations of this phenomenon, we tackle the computational challenge of exhaustively detecting minimal autocatalytic cycles (autocatalytic cores) in reaction networks, and further evaluate the impact of thermodynamic constraints on their realization under mass action kinetics. We first characterize the complexity of the detection problem by proving its NP-completeness. This justifies the use of constraint solvers to list all cores in a given reaction network, and also to group them into compatible sets, composed of cores whose stoichiometric requirements are not contradictory. Crucially, we show that the introduction of thermodynamic realism does constrain the composition of these sets. Compatibility relationships among autocatalytic cores can indeed be disrupted when the reaction kinetics obey thermodynamic consistency throughout the network. On the contrary, these constraints have no impact on the realizability of isolated cores, unless upper or lower bounds are imposed on the concentrations of the reactants. Overall, by better characterizing the conditions of autocatalysis in complex reaction systems, this work brings us a step closer to assessing the contribution of this collective chemical behavior to the emergence of natural selection in the primordial soup.

**Significance Statement:** Describing the processes behind the origin of life requires us to better understand self-amplifying dynamics in complex chemical systems. Detecting autocatalytic cycles is a critical but challenging step in this endeavor. After characterizing the computational complexity of this problem, we investigate the impact of thermodynamic realism on autocatalysis. We demonstrate that individual cycles, regardless of thermodynamic parameters, can always be activated as long as entities may occur at any required concentration. In contrast, two cycles can become mutually incompatible due to thermodynamic constraints, and will thus never run simultaneously. These results clarify the implications of physical realism for the realization of autocatalysis.

## I. INTRODUCTION

It is increasingly recognized that producing a consistent explanation for the origination of life will require us to explain how Darwinian evolution may have *gradually* emerged from a non-biological, purely physical world [1–9]. Gradually rather than suddenly, that is, without assuming that natural selection only came into play once chance alone had produced the first obvious “replicators”, displaying the same heritable variance as current organisms. Under this perspective of a smooth transition from physics to biology, natural selection is hypothesized to have been active already in the “prebiotic” soup as a driver to complexity; yet in a rudimentary and currently unrecognizable fashion.

To explore this path, autocatalysis is often taken as a plausible starting point (Box 1) [6, 10–14]. Here, more specifically, we envision autocatalytic cycles as the putative elementary components of higher level systems that may engage in “increasingly Darwinian” dynamics. In doing so, we aim at keeping the best of the two traditionally opposed approaches to the origin of life: physico-chemical realism of the metabolism-first view, and evolvability of the gene-first perspective. Beyond the specifics of terrestrial life, progressing toward an articulation of Darwinian principles with physics appears to be a prerequisite to assessing their putative relevance to other physical systems [2, 3].

We build on recent theoretical and computational developments [10, 15, 16] to systematically search for minimal autocatalytic cycles (denoted autocatalytic cores, *sensu* Blokhuis et al [15]) in reaction networks and then assess their thermodynamic consistency, i.e. the impact of thermodynamic constraints on their realization, under mass action kinetics. We first prove that finding autocatalytic cores in the network is an NP-complete problem – a question that was left open in earlier work [17, 18] – and converge with other authors in using constraint solvers as a technical solution [10, 16]. We then question whether such autocatalytic cores, defined on the sole basis of the reaction network topology, can also be realized once thermodynamic constraints are introduced. To do so, we take into account the reaction kinetics that themselves depend on the Gibbs free energies and concentrations of the reactants, and the activation barriers of the reactions. We show that regardless of these physical quantities, any potential autocatalytic core may be instantiated in some region of the concentration space as long as this space is assumed unbounded. In contrast, thermodynamic constraints do restrain compatibility relationships between autocatalytic cores and will thereby impact the dynamics in complex chemical networks.

### Box 1: Related work on autocatalysis

The present study takes place within a flourishing body of literature taking autocatalysis as a plausible primary component of proto-biotic or proto-Darwinian systems. Our model contrasts with those based on the RAF framework [19, 20] in that it follows a bottom-up approach to autocatalysis: rather than setting catalytic relationships between components of the system and randomly picked reactions, we let the reaction network generate (or not) these relationships, as formalized by Blokhuis et al [15]. Catalysis and autocatalysis then simply emerge in the reaction network as pathways involving entities that act both as reactants and products. For example, in the reactions A+C→AC, AC+B→ABC, ABC→AB + C, entity C can be simply described as a catalyst of the reaction A+B→AB.

In taking such a bottom-up angle, our framework is much related to that of several recent studies [10–12, 16, 21, 22]. Some of these have considered the implications of thermodynamic constraints and mass action kinetics on specific autocatalytic motifs [11, 12, 21]. Others have implemented tools for the exhaustive detection of autocatalysis [10, 16, 22]. Here we jointly consider these two components of the problem, i.e. exhaustive detection and thermodynamic realism.

On a more conceptual ground, we share with Baum et al [6] the view that collections of autocatalytic cycles, rather than cycles alone, might constitute the scale at which incipient heritable variations may occur.

## II. FRAMEWORK AND DEFINITIONS

We analyze networks of reversible reactions governed by mass action kinetics. This typically applies to reactions that simply consist in the association of two entities and the reciprocal dissociation (e.g. *A* + *B* ⇌ *AB*). The entities are fully defined by their composition (e.g. *A*_2_*B*_2_ is not distinct from *B*_2_*A*_2_). Given a list of entities, this rule sets the list of all possible reactions, only some of which are assumed to exist to generate a particular reaction network – this is equivalent to assuming that some reactions have an infinite activation barrier and thus have a null flow.

We can then apply the formalism of Blokhuis et al [15] to identify autocatalytic motifs in such reaction networks. Intuitively, these can be conceived as cyclic subnetworks admitting a regime where each entity has a positive net production rate.

### Definition 1.

*An autocatalytic motif is defined as a set of entities E*_*C*_ *and reactions R*_*C*_ *such that:*

- *For each reaction R* ∈ *R*_*C*_, *at least one entity on each side is in E*_*C*_.
- *There exists a vector* 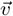 *of flows for reactions from R*_*C*_ *defining a regime where the total contribution of reactions from R*_*C*_ *is strictly positive for each entity of E*_*C*_.

Consider for instance a reaction *A* + *B* ⇌ AB, with reactants *A, B* and product *A*B. Then the stoichiometry 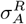 of *A* in *R* is −1 (or −2 if *A* = *B*), and 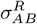 is 1. If an entity *e* does not appear in a reaction *R*, we set 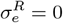.

We will note *υ*_*R*_ the flow of the reaction at a given instant, that will be positive if the association rate is larger than the dissociation rate.

Given a motif candidate *C* formed of entities *E*_*C*_ and reactions *R*_*C*_, and an entity *e* in *E*_*C*_, we define the variation of *e*’s concentration **due to** *C* as:

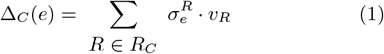

We define a *witness* as a choice of *υ*_*R*_ for each *R* ∈ *R*_*C*_, such that for each *e* ∈ *E*_*C*_ we have Δ_*C*_ (*e*) > 0. This can be formalized using linear algebra, following Blokhuis et al [15].

Indeed, if M is the stoichiometric matrix restricted to *E*_*C*_ and *R*_*C*_, then the candidate *C* is an autocatalytic motif if and only if there exists a witness vector 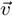 such that all coordinates of 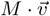 are strictly positive.

### Example 1.

*We illustrate the definition in Figure 1 with a simple formose-like cycle comprising three entities A*_2_, *A*_3_ *and A*_4_ *and using entity A*_1_ *as food, with detailed explanations provided in the caption*.

**Figure 1.**
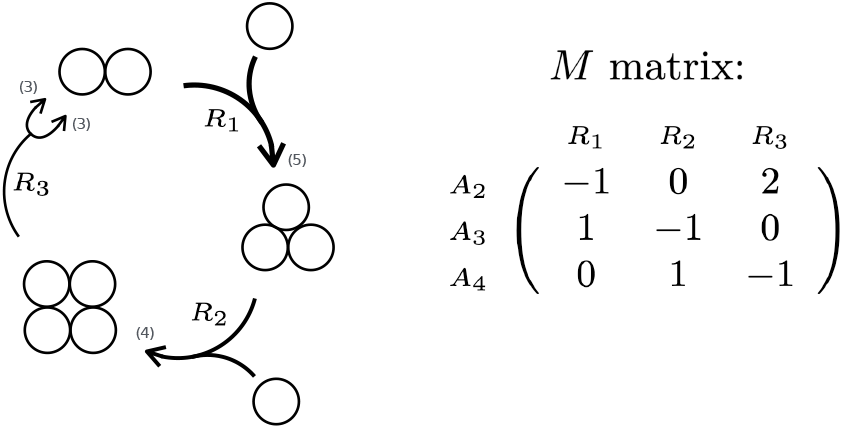
Schematic view of a formose-like autocatalytic motif. Entities *A*_2_, *A*_3_ and *A*_4_ are part of the motif while entity *A*_1_ serves as food. The arrows indicate the net direction of each reaction, while the line width indicates their respective flows, that must be decreasing from reactions *R*_1_ to *R*_3_ for the cycle to run. As an example, the flows values (indicated in brackets) would produce a net increase of 1 of each entity. The right panel shows the corresponding stoichiometric matrix *M*. Given the represented flow vector 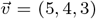, we obtain 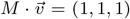, showing that 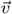 is indeed a witness.

Entities appearing in *R*_*C*_ that are not part of *E*_*C*_ will be called either “food” if they are consumed or “waste” if they are produced by a reaction of *R*_*C*_, taking into account the sign of the witness vector that indicates the direction of reactions. Notice that an entity may simultaneously appear as food and waste in an autocatalytic motif.

Formally, a motif is said to *contain* another one if it includes all its entities and reactions. In contrast, a motif is *minimal* if it does not contain any other, in which case it corresponds to the “autocatalytic core” from Blokhuis et al [15] and to the stoichiometric autocatalysis of Gagrani et al [16]. The present study focuses on such minimal motifs, that we denote *Potential* Autocatalytic Cores (PACs), to emphasize that they are defined on the sole basis of the reaction network topology, so that their realizability under thermodynamic constraints remains to be assessed.

### Definition 2.

*A Potential Autocatalytic Core (PAC) is defined as a minimal autocatalytic motif, whose identification relies solely on the reaction network topology as formalized by the stoichiometric matrix*.

### Example 2.

*The autocatalytic motif shown in Figure 1 is minimal, and hence constitutes a PAC*.

Notice that minimality implies that a PAC witness cannot contain null reaction flows: if this were the case, the corresponding reaction could be removed without altering autocatalysis, which contradicts the minimality criterion. Furthermore, it is shown in Blokhuis et al [15] that in a PAC, each entity is the reactant of a unique reaction, and each reaction has a unique entity of the PAC as reactant (other reactants being food). This implies that the direction of reactions is consistent across all witnesses of a given PAC: flipping the direction of one reaction would impose flipping all the others. Therefore, each reaction of the PAC has a unique possible net direction that will be compatible with all its witnesses flow vectors. In Appendix 1, we provide a formal proof of this property that was put forward in Blokhuis et al [15, SI].

## III. DETECTING POTENTIAL AUTOCATALYTIC CORES

Our goal is to enumerate all PACs in a reaction network. To this end, we first assess the complexity of this problem, in order to determine which computational tools are required for its resolution.

### A. NP-completeness proof

Enumerating all PACs in a reaction network involves sequentially solving problems of the type: “Is there a PAC in the system besides those previously found?”. We will show that a particular variant of this question is NP-complete. Namely, we will prove the NP-completeness of deciding whether a PAC exists that contains an entity *A* and takes food from a given subset *F*.

In this section, for simplicity, we will relax any compositionality constraint on reactions so that letters like *A, B, E* … will be shorthand for any kind of entity. Yet it would be straightforward (but less readable) to extend the construction to a strictly compositional framework.

Notably, the complexity of the autocatalysis detection problem has previously been considered by Andersen et al but from a different angle. These authors have specifically shown that the following problem is NP-complete: considering a reaction network that contains a known autocatalytic core, can its resources be produced by the network? The difficulty of finding all autocatalytic cores in a reaction network, that we tackle here, has thus not yet been addressed.

In the framework of Blokhuis et al [15], it is easy to check whether a proposed set of entities and reactions constitutes an autocatalytic core. Indeed, thanks to the linear algebra formulation summarized in Section II, this problem is solved in polynomial time by Linear Programming. As will be shown, the difficulty rather lies in finding an autocatalytic core in a reaction network, among exponentially many possible candidates.

Formally, let PAC-DETECTION be the following algorithmic problem:

#### Definition 3

(PAC-DETECTION problem).

***INPUT***: *A reaction system defined by entities* (*E*) *and reactions* (*R*), *a target A* ∈ *E, and a set of allowed foods F* ⊂ *E*.

***OUTPUT***: *Is there a PAC containing A and using only foods from F ?*

#### Theorem 1.

*PAC-DETECTION is NP-complete*.

We provide here a brief description of the proof that is fully described in Appendix 2. Because a PAC candidate can be tested in polynomial time, PAC-DETECTION is in NP. It remains to be shown that it is NP-hard. To this end, we reduce from the well-known NP-complete problem SAT [23]. An instance of SAT asks whether an input formula on *n* boolean variables *x*_1_, …, *x*_*n*_ is satisfiable, via a suitable assignation of variables with true/false values.

To perform the reduction, we associate each such formula *φ* with a reaction system *S*_*φ*_, of size polynomial in *φ*, with a specified target entity *A* and a food set *F. R*eactions in *S*_*φ*_ are designed to mirror the structure of *φ*, ensuring that a PAC of the wanted form exists if and only if the formula is satisfiable. The only possible such PACs will actually directly encode satisfying assignments for *φ*.

This shows that PAC-DETECTION is NP-hard: a polynomial-time algorithm for PAC-DETECTION would yield a polynomial-time algorithm for SAT, via this reduction. We can conclude that PAC-DETECTION is NP-complete, since it is also in NP. Notice that our proof assumes a particular shape for constraints on the searched PAC (containing an entity *A* and using only foods from a set *F*). Assessing whether an unconstrained variant is NP-complete as well remains an open problem.

### B. Implementation

The above NP-completeness result justifies the use of an SMT Solver such as Z3 to enumerate all PACs. We thus implemented this approach in C**++** in the EmergeNS software [24] (more generally designed to simulate the dynamics of complex physicochemical systems and down the line to trace the physical emergence of natural selection). In practice, in a reaction system defined in EmergeNS, we ask the Z3 solver to find PAC candidates and to assess, for each candidate, the existence of a PAC witness, i.e. a reaction flow vector 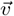 yielding strictly positive net production rates for all entities of the candidate. We then remove the detected PAC from the search space and repeat the process until no new PAC is found.

As remarked in Section III A, verifying whether a given candidate is indeed a PAC (by assessing the existence of a PAC witness) is a linear programming problem, and can thus be achieved efficiently, i.e. in polynomial time. The need for Z3 comes from the search for PAC candidates in an exponential space of possible subsets of entities and reactions.

## IV. PAC CONSISTENCY UNDER THERMODYNAMIC CONSTRAINTS

The kinetics of a reaction network must obey the second law of thermodynamics, a constraint that is not considered in the PAC definition. Indeed, this definition solely relies on the existence of a witness 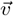 of reaction flows, that may or may not be compatible with thermodynamic constraints.

More precisely, once association and dissociation constants are derived from free energies and activation barriers, they cannot be freely chosen. To clarify these constraints, recall that each entity *e* is associated with a chemical potential *µ*, which is the sum of its molar Gibbs free energy (or standard chemical potential) *G* and the logarithm of its activity (that we identify here to its concentration [*e*], hence placing ourselves in the ideal solution regime)^1^:

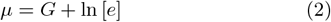

In addition, the flow of a reaction *R*, denoted *υ*_*R*_, depends on the concentrations (i.e. activities) of its entities, under mass action kinetics:

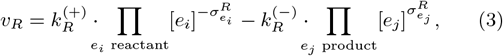

Where 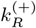 and 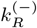 are respectively the forward and backward kinetic rate constants and 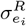 is the stoichiometry of 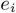 in reaction *R* (negative for reactants, hence explaining the presence of a minus sign in the above formula). The key point, to enforce the second law under mass action kinetics, is to relate the rate constants to Gibbs free energies using the local detailed balance condition (also known as Eyring’s formula):

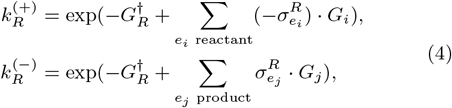

where 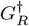 is the free energy of the intermediate state that induces an activation barrier. Injecting Equation (4) in Equation (3) leads to the following formulation that will be further used below:

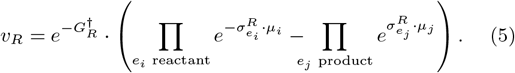

The sign of *υ*_*R*_ is the same as that of the affinity of the reaction 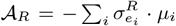, expressing how a positive rate can be achieved. *R*ecall, however, that positive flows for all reactions of a PAC will not necessarily entail autocatalysis, as we aim more specifically for flows balanced in such a way that each entity has a net positive production rate (Definition 1). The question of addressing the thermodynamic consistency of PACs can thus be reformulated as follows: can a PAC flow witness 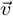 be realized through a concentration vector 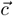 of all entities of the system? If such a vector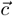 exists, the PAC will be considered a thermodynamically Consistent Autocatalytic Core (CAC). Here, “thermodynamic consistency” refers to mass action kinetics, with rate constants derived from Gibbs free energies and activation barriers.

### Definition 4.

*A Consistent Autocatalytic Core (CAC) is a PAC for which there exists a concentration vector* 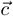 *yielding reaction flows forming a PAC witness*.

The concentration vector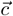 will then be called a *CAC witness*. Here, we reach the second key result of the present study, namely, that one can always find such a CAC witness – i.e. that a single PAC is always thermodynamically consistent – as long as the concentration space is unbounded.

### Theorem 2.

*Let* 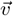 *be a PAC witness (or any flow vector compatible with the directions of the PAC reactions). Then there exists* λ > 0 *and a concentration vector*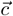 *such that* 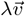 *is the flow vector induced by* 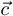.

The general proof of this theorem is given in Appendix 3, with an example provided in Appendix 3 a. It should be noted that Theorem 2 actually has a broader scope than the problem specifically addressed in this section, since it demonstrates that *any* flow vector 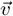 matching the directions of the PAC reactions can be realized in the concentration space up to some proportionality factor λ > 0. Notably, this holds for any values of activation barriers and Gibbs free energies, and even when food and waste concentrations are fixed to any arbitrary values.

This goes beyond the well-established result, based on affinity, that a positive flow can be achieved for any reaction by adjusting concentrations. Here, the difficulty is to aim for a fixed vector 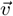 of rates, up to some coefficient λ shared among all reactions of the PAC.

We deduce the following corollary from Theorem 2:

**Corollary 1**. *Any PAC is a CAC*.

*Proof of Corollary 1*. Consider a PAC formed of entities (*e*_1_, …, *e*_*n*_) and reactions (*R*_1_, …, *R*_*n*_) (recall that according to Blokhuis et al [15], the minimality of a PAC implies that it contains as many entities as reactions). Let 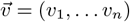 be a PAC witness. Since the inequalities to be satisfied by a PAC witness are all linear with respect to the coordinates of 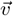 (see Equation (1)), for all λ > 0, 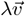 is a PAC witness as well. Applying Theorem 2 gives us the existence of some λ > 0 and a concentration vector 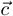 yielding flows 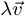, which is a PAC witness. Thus, 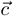 is a CAC witness, and the arbitrary PAC we started with is indeed a CAC.

## V. COMPATIBILITY AMONG AUTOCATALYTIC CORES

In this section, we investigate whether thermodynamic constraints affect compatibility relationships among cores. To do so, we first analyze compatibility among PACs, that is, we identify sets of cores that are found compatible on the basis of the reaction network topology alone, hereafter called “multiPACs”. A set of PACs is a multiPAC if there exists a common witness 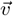 of reaction flows allowing all the PACs of the set to run simultaneously. In the framework of Gagrani et al [16], this corresponds to a nonempty intersection of the flow-productive cones for the different autocatalytic cores considered. Here, instead of computing explicit intersections, we will use the Z3 solver to identify nonempty intersections, and directly ask for a single vector 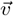 witnessing the different PACs simultaneously. To do so, we simply concatenate the requirements already defined for each individual PAC. Let us emphasize that a multiPAC witness does not guarantee global production of each entity when reactions from all PACs are jointly considered (this can be seen for example in Figure 4, where entity *e*_4_ has a positive production rate within each PAC, although it is consumed overall).

Interestingly, we note that incompatibilities between PACs may occur for various reasons. The simplest case, illustrated in Figure 2, is when a reaction is shared between two PACs, but in opposite directions. This obviously prevents the existence of a common flow vector witness 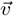. Yet more subtle cases were also obtained with our software using randomly generated reaction networks. One example is described in Figure 3. Generally speaking, incompatibilities among PACs occur because of contradictory flow requirements, that is, when one PAC requires a reaction *R*_1_ to run faster than a reaction *R*_2_, while a second PAC requires the opposite.

**Figure 2.**
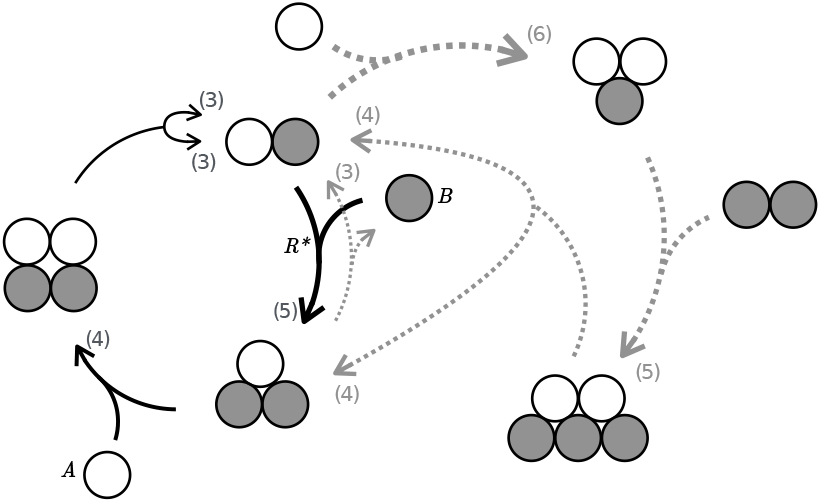
A simple example of two incompatible PACs. Ignoring foods and wastes and identifying A to empty circles and B to filled circles, the PAC with solid black arrows is *AB* → *AB*_2_ → *A*_2_*B*_2_ → *AB* + AB and the PAC with gray dotted arrows is *AB*_2_ → AB → *A*_2_*B* → *A*_2_*B*_3_ → *AB* + *AB*_2_. Numbers in brackets indicate the flows of PAC reactions allowing for a (local) net production of 1 of all their entities. The two PACs share the reaction labeled *R*^⋆^, but require it to run in opposite directions.

**Figure 3.**
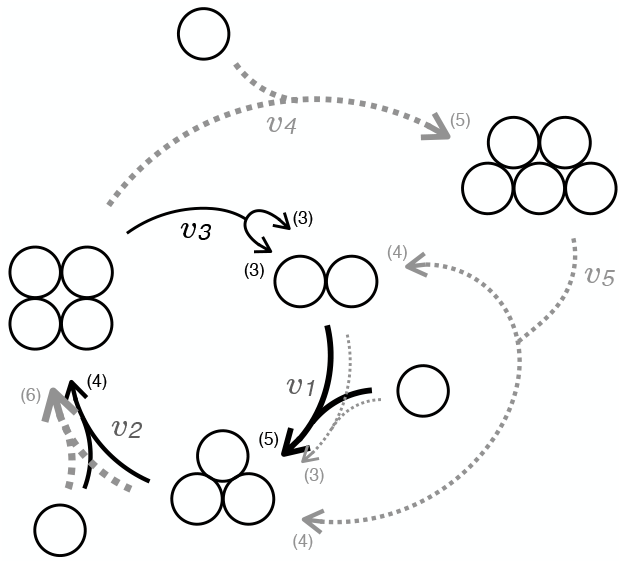
A more subtle example of two incompatible PACs, sharing two reactions indexed with flows v_1_ and v_2_. Flows allowing for unitary production of PAC entities are shown in brackets for both PACs. The inner PAC depicted with solid black arrows (*A*_2_ → *A*_3_ → *A*_4_ → *A*_2_ + *A*_2_) imposes the flow inequalities *υ*_1_ > *υ*_2_ > *υ*_3_ > *υ*_1_/2, while the gray dotted outer PAC (*A*_2_ → *A*_3_ → *A*_4_ → *A*_5_ → *A*_2_ + *A*_3_) imposes inequalities *υ*_1_ + *υ*_5_ > v_2_ > *υ*_4_ > *υ*_5_ > *υ*_1_. This yields contradictory requirements: *υ*_1_ > *υ*_2_ for the first PAC and *υ*_1_ < *υ*_2_ for the second.

To investigate the impact of thermodynamic constraints on compatibility relationships among cores, we now assess whether pairs of compatible PACs, witnessed by a common flow vector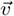, also constitute pairs of compatible CACs, witnessed by a common concentration vector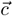. As before, addressing this question using the SMT solver is straightforward: it is enough to concatenate the lists of constraints required for each of the CACs under study, and to ask if a 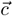 vector exists that simultaneously satisfies them all.

This analysis reveals that compatibility among cores inferred solely from the reaction network topology may overlook thermodynamic inconsistencies. Indeed, we find several instances of two compatible PACs making incompatible CACs. We give an example of such a behavior in Figure 4 and provide a formal proof that these two CAC are incompatible in Box 2.

### Box 2: Thermodynamic incompatibility between CACs

On the example shown in Figure 4, we can first show that the two PACs are compatible, since we can choose a set of reaction flows that witnesses both of them simultaneously. For instance setting 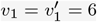, v_2_ = 5, v_3_ = 4, 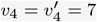 leads to a positive production rate of each entity by each PAC. More generally, this criterion is achieved for both PACs simultaneously when the following inequalities are satisfied: *υ*_3_ < *υ*_2_ < *υ*_1_ < v_4_ < 2v_3_ and 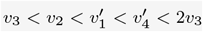.

**Figure 4.**
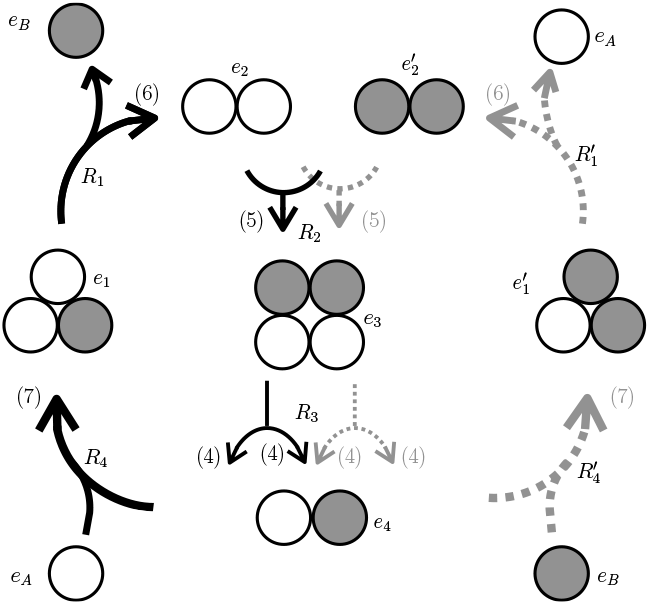
Two topologically compatible but thermodynamically incompatible autocatalytic cores. The two PACS (*{R*_1_, *R*_2_, *R*_3_, *R*_4_*}* and 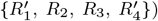 have two reactions in common. One can check that both cores run simultaneously with flows 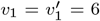, *υ*_2_ = 5, *υ*_3_ = 4 and 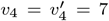 which allows for a production rate of 1 for all entities belonging to a PAC. However, as detailed in Box 2, it can be shown that these cores cannot be instantiated simultaneously in the concentration space, i.e. they form a multi-PAC but not a multiCAC.

We will show that these inequalities are not thermody-namically achievable, i.e. that they lead to contradictory requirements in the concentration space.

Notice that the inequalities imply that all *υ*_i_ are strictly positive, because *υ*_3_ < 2*υ*_3_ entails *υ*_3_ > 0, and all other *υ*_i_ are larger than *υ*_3_. As proven below, this sign constraint alone is not satisfiable: not all reactions can flow in the wanted direction.

We denote 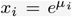 the exponential of the chemical potential of entity *e*_i_ (indexed as in Fig. 4), and 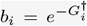 where 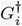 is with respect to reaction *R*_i_ (similarly for 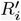). Reaction flows can thus be written as follows:

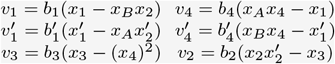

From the positivity of all flows, we get:

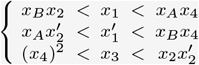

From the first line we deduce *x*_B_*x*_2_/*x*_4_ < *x*_A_, and reinjecting in the second line we obtain 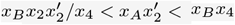. This simplifies to 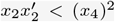, which contradicts the condition of the third line. It should be noted that this proof is valid regardless of the activation barriers values, since the contradiction stems from the signs of the flows, while activation barriers only affect their amplitudes.

## VI. DISCUSSION

Working toward the long-term goal of an explicit grounding of Darwinian dynamics into physical processes, we addressed in this study the implications of thermodynamic constraints on the existence and detection of autocatalytic cores given a reaction network. Our analysis builds on recent theoretical progresses made on the formalization of autocatalysis on the sole basis of the reaction network topology [15]. Under this definition, we show that the exhaustive detection of autocatalysis is an NP-complete problem. This finding fully justifies the use of constraint solvers (e.g. SMT, Integer Programming) toward which we converge with others [10, 16].

The constraints imposed by free energies and activation barriers can always be compensated by adjusting concentrations, thereby allowing any minimal autocatalytic cycle to also be thermodynamically consistent under mass action kinetics. In other words, the list of autocatalytic cores in a reaction network remains unaffected by these physical constraints, as long as concentrations are not limited by upper or lower bounds. However, as shown in Appendix 4, it should be noted that heterogeneity in free energies and activation barriers do restrict the volume of the concentration space where a core can effectively run.

These conclusions on isolated cores do not readily apply to combinations of cores. Indeed, thermodynamic realism does restrict the list of mutually compatible cores, even in an unlimited concentration space, so that topologically compatible cores can turn out incompatible. Incompatibilities between two autocatalytic cores can therefore stem from two distinct sources, namely the topology of the reaction network (PAC-incompatibility) and irreconcilable demands on concentrations (CAC-incompatibility).

A stimulating next step will be to investigate the implications of autocatalysis on the system’s dynamics through time and space. Indeed, the existence of a CAC witness says little of the actual trajectories occurring in the concentration space, and on the influence of autocatalytic cores on these trajectories. Among the many autocatalytic cores that are thermodynamically achievable in a given system, which ones are actually encountered from a given starting point in the concentration space? Which ones are short-lived, long-lived, or keep running once the system has reached a steady state? Which ones run in the vicinity of this steady state, and perhaps contribute to drive the system in its direction? And finally, could autocatalysis contribute to generating more than one steady state in the concentration space? We anticipate that such multistability could enable a primordial form of heritable variation, paving the way to nascent Darwinian dynamics.

## ACKNOWLEDGMENTS

We are very grateful to Nicolas Lartillot for his insightful contribution in setting up our modeling approach, and to Benjamin Kuperberg for sharing his open source GUI library OrganicUI. We also thank *R*omain Yvinec, Olivier *R*ivoire, Yann Sakref and Iris Magniez--Papillon for fruitful discussions, as well as Arnaud Mary, two anonymous reviewers and the associate editor for constructive comments on earlier versions of the manuscript. Thomas Kosc was supported by grant AN*R*-17-CE02-0021-01 to Sylvain Charlat.

## APPENDIX

### 1. Uniqueness of reaction directions in PAC witnesses

*R*ecall that a PAC in our framework is defined as an autocatalytic core from Blokhuis et al [15]. We will use some known properties of such cores:

#### Lemma 1

(from Blokhuis et al SI [15]). *Any autocatalytic core is of the following form:*

1. *entities e*_1_, …, *e*_*n*_,
2. *reactions R*_1_, …, *R*_*n*_ *(i*.*e. same number as entities)*,
3. *for each i* ∈ [1, *n*], *the sole reactant of R*_*i*_ *among e*_1_, …, *e*_*n*_ *is e*_*i*_,
4. *the directed graph G with vertices* [1, *n*] *and edges* {(*i, j*) | *e*_*j*_ *appears as product in R*_*i*_} *is strongly connected*.

We show that this characterization implies that the direction of any reaction cannot be reversed in PAC witnesses. This was mentioned in [15, SI], with the sentence: “we choose the convention that any productive vector *γ* is positive, up to taking the opposite for some columns of M”, but we will prove this here for completeness.

First of all, if a reaction *R*_*i*_ has two distinct products *e*_*j*_, *e*_*k*_, then the reverse reaction has two distinct reactants among the PAC entities, and is therefore incompatible with the third item of Lemma 1, so such a reaction cannot be reversed.

We now assume for contradiction that some reaction may be reversed in a PAC witness. Let us call 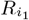 this reaction, with original reactant 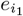 and product 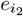, with *i*_1_ ≠ *i*_2_. Our assumption is that there exists another PAC witness for the same PAC, where 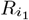 runs in reverse, that is with reactant 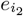 and product 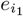. Since 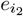can only appear once as reactant according to Lemma 1, this forces us to reverse reaction 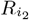 as well. The former product 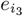 of this reaction becomes a reactant, forcing us to reverse reaction 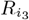. Continuing this process, one of these events will eventually occur:

1. We encounter a reaction 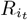 having two distinct products. Since we cannot reverse it, the entity 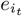 will appear twice as reactant, contradicting Lemma 1.
2. We encounter a reaction 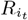 whose unique product is 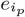 with *p* < t, i.e. 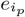 was already encountered in the process. Then, by the fourth item of Lemma 1, we must have t = *n* and *p* = 1. That is, this process has reached all reactions and now loops back to the starting point.

Therefore, all that remains is to show that switching the direction of all reactions leads to a contradiction as well. Since all reactions have a unique product among the entities of the PAC, we are here in the case of a type I core in the terminology of [15]. We can renumber reactions so that in the original PAC, the unique product of reaction *R*_*i*_ is *e*_*i*+1_ for *i* < *n*, and *e*_1_ for *i* = *n*. Let *a*_*i*_ be the absolute value of the stoichiometric coefficient of reactant *e*_*i*_ in *R*_*i*_, and b_*i*_ be the stoichiometric coefficient of the product of this reaction (*e*_*i*+1_ or *e*_1_ for *i* = *n*).

Let 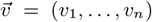 be a PAC witness consistent with the original directions of reactions of the PAC, with *υ*_*i*_ > 0 for all *i* ∈ [1, *n*]. *R*ecall that as mentioned in Section II, no coordinate of 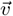 can be null, as the corresponding reaction could be removed, contradicting minimality of the PAC. What we aim to show is that there exists no other PAC witness 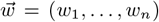, with w_*i*_ < 0 for each *i*, that would correspond to reversing the direction of all reactions.

The fact that 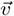 is a PAC witness is expressed by the following inequalities:

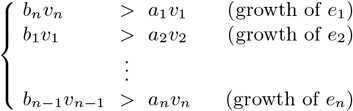

Multiplying these inequalities all together and simplifying by *υ*_1_*υ*_2_ … *υ*_*n*_ (which is strictly positive) yields b_1_b_2_ … b_*n*_ > *a*_1_*a*_2_ … *a*_*n*_.

This highlights the weight-asymmetry characterizing Type I cores as stated in [15, SI], with an additional constraint: the product of stoichiometric coefficients associated with reaction products is not only different from the product of absolute values of stoichiometric coefficients associated with reactants, but it must be strictly larger. This is enough to show that reversing the directions of all reactions is not possible, since this inequality would then be in the wrong direction. This completes the proof that the sign of flows in PAC witnesses cannot be altered.

#### 2. Proof of Theorem 1: NP-completeness

This section is devoted to the proof of Theorem 1.

First, by the above remark, if a candidate PAC is given then it can be checked in polynomial time; thus PAC-DETECTION is in NP. To achieve the proof, it remains to be shown that it is NP-hard. We do so by reducing from the classical NP-complete problem SAT, that is, by translating the SAT problem into PAC-DETECTION.

An instance of SAT is a boolean formula *φ* on *n* variables *x*_1_, *x*_2_, …, *x*_*n*_. We will call literal a variable *x*_*i*_ or its negation 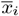. The formula *φ* is a conjunction of k clauses, i.e. of the form *φ* = ∧_1≤*j*≤k_ *c*_*j*_, where each clause c_*j*_ is a disjunction of literals. The formula *φ* is satisfiable if there exists a valuation setting a boolean value for each *x*_*i*_ (and the opposite for 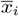), that allows *φ* to evaluate to *true*. The SAT problem, that is, asking whether an input formula *φ* is satisfiable, is known to be NP-complete [23].

Let us now encode SAT into PAC-DETECTION. Given an instance of SAT, i.e. a formula *φ* as above, we want to design a reaction system *S*_*φ*_ = (*E, R*), together with a target entity *A* ∈ *E* and a food set *F* ⊂ *E*, such that there is a PAC satisfying this instance of PAC-DETECTION if and only if *φ* is satisfiable. This would mean that a (hypothetical) polynomial-time algorithm for PAC-DETECTION would yield a polynomial-time algorithm for SAT, provided the reduction can be done in polynomial time, and produces a reaction system of polynomial size, which will be ensured here.

We choose as entities the set:

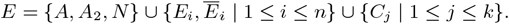

This means we have 3 + 2*n* +*k* entities in the system. Note that we use capital letters to distinguish entities from the variables and clauses from *φ* to which they refer.

For reactions, we choose the following set, where *L*_*i*_ always stands for either *E*_*i*_ or *Ē*_*i*_, e.g. the first line of the list of reactions shown below means that both reactions *A* ⇌ *E*_1_ + N and *A* ⇌ *Ē*_1_ + N are present:

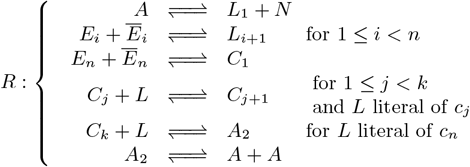

Finally, the target is *A* and the allowed foods will be *F* = {*E*_*i*_, *Ē*_*i*_ | 1 ≤ *i* ≤ *n*}.

The idea is that a valuation witnessing satisfiability of *φ* will correspond to a PAC of the form:

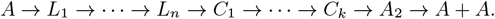

The crux of the construction is that reactions using *C*_*j*_ as reactant need one of the literals of c_*j*_ as food, that will be interpreted as the literal validating the clause *c*_*j*_. In consequence, this literal must be a food, and cannot appear as one of the entities of the PAC. So the segment *L*_1_ → … → *L*_*n*_ appearing in the PAC has to be formed exactly of the literals that are put to false in the valuation.

The entity N is used to force the direction of the reaction *A* → *L*_1_ in the PAC: since it can only be used as waste and not as food, the reaction cannot go in the opposite direction. From there, any PAC containing *A* and using only foods from *F* must be of the above form, and witnesses a valuation satisfying *φ*.

##### Example 3.

^2^ *Consider φ given by c*_1_ = *x*_1_ ∨*x*_2_, 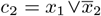, *and –*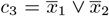. *Then the only correct valuation is the one setting x*_1_ *to true and x*_2_ *to false. This is witnessed in the system S*_*φ*_ *by the PAC A* → Ē_1_ → *E*_2_ → *C*_1_ → *C*_2_ → *C*_3_ → *A*_2_ → *A* + *A, using E*_1_ *and Ē*_2_ *as foods (and* N *as waste)*.

*If we add a clause* 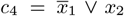 *to φ, the formula becomes unsatisfiable and accordingly, the corresponding system S*_*φ*_ *does not contain a PAC satisfying the constraints*.

This achieves the proof that PAC-DETECTION is NP-complete.

#### 3. Proof that all PACs are CACs

In the following, to simplify notations, reactions will be oriented in accordance to the PAC, i.e. all coordinates of the PAC witness will be positive. A “positive flow vector” is a vector where all coordinates are positive.

##### a. An example

###### Example 4.

*Let us first instantiate the proof with the PAC mentioned earlier, given by the following reactions (ignoring foods)* 2*e*_1_ → *e*_2_ → *e*_3_ → *e*_1_ + *e*_2_, *depicted in Figure 5*.

**Figure 5.**
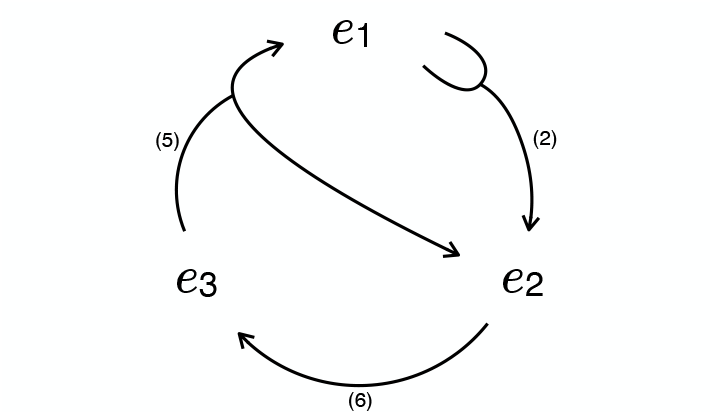
A PAC example of size 3 to give a first intuition of the proof.

*A PAC witness is* 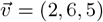, *ensuring a production rate of* 1 *for each entity. Using Equation* (5), *we now look for a concentration vector to realize this PAC witness with the following flows:*

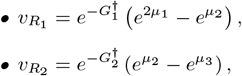

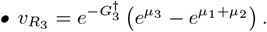

*We will instead aim at realizing the flow vector* 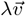*for some* λ > 0, *where* 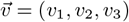 *is the original PAC witness, e*.*g*. 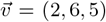. *Letting* 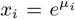 *and* 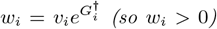, *the equation system can be written conveniently:*

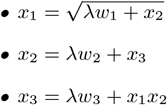

*Substituting x*_2_ *we obtain* 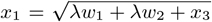, *and finally substituting x*_1_ *as well we obtain:*

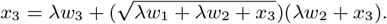

*Let us note g*(λ, *x*_3_) *the above right-hand side expression, so that the equation becomes x*_3_ = *g*(λ, *x*_3_). *Let us define h*(λ, *x*_3_) = *g*(λ, *x*_3_) − *x*_3_, *so that we aim at h*(λ, *x*_3_) = 0. *We will show that such a solution exists using the intermediate value theorem*.

*For x*_3_ = 0 *and any* λ > 0, *we have* 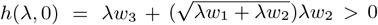 *Let us choose any a* ∈ (0, 1). *For x*_3_ = *a and* λ = 0, *we have h*(0, *a*) = *a*^3/2^ − *a* < 0. *Because h is continuous there exists ε* > 0 *such that for all* λ ∈ (0, *ε*), *h*(λ, *a*) < 0. *Let us choose* λ ∈ (0, *ε*), *we know thanks to the intermediate value theorem (on x*_3_ *with this fixed value of* λ*) that there exists x*_3_ ∈ (0, *a*) *such that h*(λ, *x*_3_) = 0. *From this x*_3_ *we can compute* 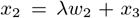, *and* 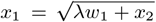.

*We have obtained a solution to our system by an appropriate choice of* λ, *x*_1_, *x*_2_, *x*_3_, *thereby witnessing that the PAC is a CAC via Corollary 1*.

##### b. General proof

We aim to prove Theorem 2 from Section IV by generalizing the proof exposed in the previous example to any PAC.

**Notations**

Let 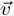 be a positive flow vector, we want to show that there exists λ > 0 and a concentration vector 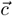 such that 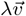 is the flow vector induced by 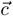. We will even show that this can be attained for any fixed concentration values of food and waste entities.

Let 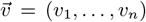 be the target flow vector, where all coordinates are strictly positive. Here *n* is the size of the PAC, i.e. both the number of entities *e*_1_, …, *e*_*n*_, and reactions *R*_1_, …, *R*_*n*_, where for each *i, e*_*i*_ is the sole reactant of *R*_*i*_ among entities of the PAC, see discussion in [15].

For an entity *e*_*i*_ (1 ≤ *i* ≤ *n*) with Gibbs free energy 𝒢_*i*_, recall that its chemical potential is given by *µ*_*i*_ = 𝒢_*i*_ +ln ([*e*_*i*_]). We will be interested here in the exponential of this potential, a variable that we denote 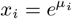.

Notice that *x*_*i*_ can be freely adjusted to any strictly positive value by choosing the appropriate concentration [*e*_*i*_], so we can turn the problem into that of finding a solution vector 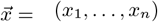. Analogs of components of 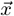 associated to food and waste entities are fixed to 1 for simplicity. Note that this does not involve chemostatting, but simply that a particular solution (a CAC witness) of a set of polynomial equations is searched, and that it is enough to fix food and waste variables to 1 to find such a solution. This does not mean that we impose a constant concentration of those entities over time. The proof can easily be adapted to any given values, but this will save us some notations, as food and waste entities can now be ignored in the computation of the reaction flows. One can refer to Appendix 4 for an analysis of the impact of food and waste potentials on the feasibility of PACs.

Let us recall Equation (5) giving the flow of a reaction in terms of chemical potentials:

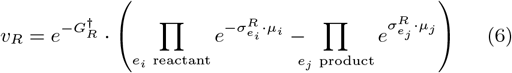

In our case, given that *e*_*i*_ is the sole reactant of *R*_*i*_ (apart from possible foods), this can be rewritten for any *i* ∈ [1, *n*]:

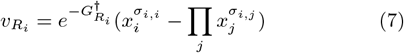

Where *σ*_*i,i*_ is the stoichiometry of the reactant *e*_*i*_ in *R*_*i*_, and *σ*_*i,j*_ is the stoichiometry of product *e*_*j*_ in *R*_*i*_. It should be noted here that for simplicity we change the convention regarding the stoichiometry of reactants, so that all parameters are positive, including *σ*_*i,i*_.

Let us note 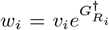. The fact that we aim for 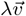 to be the flow vector induced by 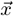 means that for all *i*∈ [1, *n*], we aim for 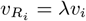, so we must have 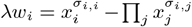 and finally:

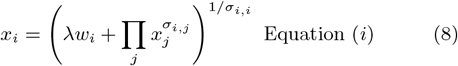

Notice that *σ*_*i,i*_ > 1 corresponds to “inverted forks”, where several *e*_*i*_ must be used as reactants. This can indeed occur in a PAC, as witnessed by Example 4.

###### Variable elimination

Our goal is to find λ > 0 such that this system has a solution (*x*_1_, …, *x*_*n*_) with each *x*_*i*_ > 0. We will do so by iteratively reducing the number of variables and equations. First, we perform the following operation: as long as there exists an equation of the form *x*_*i*_ = *A*_*i*_, where *A*_*i*_ is an expression not depending on *x*_*i*_, we replace *x*_*i*_ by *A*_*i*_ in all other equations, and remove Equation (*i*).

We will end up with a list of equations of the form *x*_*i*_ = *g*_*i*_(λ, *x*_1_, …, *x*_*n*_) where *i* ranges over some subset of [1, *n*], and where *g*_*i*_ is an expression depending on some of its variables including *x*_*i*_, and generated by the following grammar:

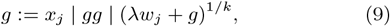

where *j* ranges over [1, *n*], and k is a strictly positive integer. There are as many remaining equations as remaining variables. Without loss of generality, we assume that the remaining variables are *x*_1_, …, *x*_*p*_, where *p* ≤ *n* is the number of remaining equations.

###### System rewriting as graph transformations

Before tackling the resolution of the system per se, we relate some properties of the remaining equations to the topology of the PAC we started with. We associate the PAC with a graph 𝒢, whose vertices are entities *e*_1_, … *e*_*n*_, and in which there is an edge *e*_*i*_ → *e*_*j*_ if *e*_*j*_ is among the products of the reaction using *e*_*i*_ as reactant.

Substituting variable *x*_*i*_ in the system according to Equation (*i*) amounts to removing vertex *e*_*i*_ and contracting all edges originating in this vertex, as shown in Figure 6 with *i* = 5:

**Figure 6.**
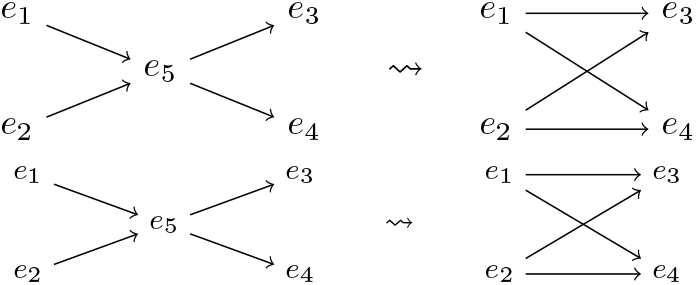
The effect of substituting the equation *x*_5_ = λ*w*_5_ + *x*_3_*x*_4_ on the graph 𝒢.

This operation can only be performed if there is no selfloop on *e*_*i*_ in the current graph. Therefore, the case where we can end up with only one equation on an entity *x*_*i*_ (i.e. the case *p* = 1) corresponds to the fact that the graph 𝒢 \ {*x*_1_} is acyclic.

We give in Figure 7 an example where this does not happen, no matter in which order the substitutions are done. This example corresponds to the Type V pattern from [15].

**Figure 7.**
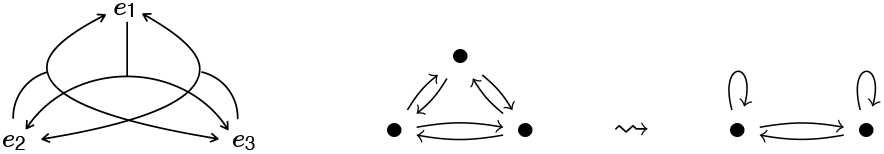
A 3-entity PAC, and its graph 𝒢. After performing one substitution step, all nodes have self-loops and the substitution sequence must end, with equations of the form e.g. *x*_1_ = λ*w*_1_ + (λw_3_ + *x*_1_*x*_2_)*x*_2_.

###### Shape of the reduced system

Let us now turn to the following lemma, which gives some properties of the expressions *g*_*i*_ reached by this process.

###### Lemma 2.

*For all* 1 ≤ *i* ≤ *p*,

1. *g*_*i*_ *is always of the form* 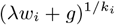,
2. *for* λ = 0, 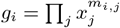 *with m*_*i,j*_ ≥ 0 *for all j, and*
3. *if only one equation remains (p* = 1*), m*_1,1_ > 1,
4. *otherwise if p* > 1, *then for all i* ∈ [1, *p*], *we have m*_*i,i*_ = 1 *and m*_*i,j*_ > 0 *for some j* ≠ *i*.

*Proof*. The first item is a direct consequence of the operation we performed, starting from Equation (*i*).

The general shape of the second item is guaranteed by construction.

We now show item 2.1. Let us first give an interpretation for exponents *m*_*i,j*_. Consider Equation (*i*) when λ =0:

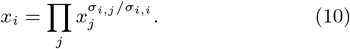

Recall that we have defined *σ*_*i,i*_ > 0 even though entity *e*_*i*_ is a reactant in reaction *R*_*i*_. Exponents in the equation system for λ = 0 simply express the fact that in reaction *R*_*i*_ and starting from one unit of entity *e*_*i*_, for all *j* one can produce *σ*_*i,j*_/*σ*_*i,i*_ units of entity *e*_*j*_. The substitution algorithm is equivalent to consuming all available *e*_*j*_ via reaction *R*_*j*_, with exponents of the products keeping track of their quantity. By iteration, if one started with entity *e*_*i*_ and could remove all reactions from the system (in our notation that would mean *p* = 1), one should recover that *e*_*i*_ can self-amplify through the reactions of the core since it allows for autocatalysis of *e*_*i*_. This intuitively explains why we expect *m*_1,1_ > 1. For illustration in Example 4, the core allows the producing of 3/2 units of entity *e*_3_ starting from one unit of it.

Let us now give a formal proof of the fact that *m*_1,1_ > 1 when *p* = 1, and let us note *m* = *m*_1,1_ for concision. The argument is similar as the proof from Appendix 1. The sequence of substitutions leading to the expression *g*_1_ amounts to describing a positive flow vector 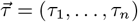 in the following way: the flow of reaction *R*_1_ is set to τ_1_ = 1, and then all available entities *e*_*j*_ other than *e*_1_ are entirely consumed and substituted with the products of reaction *R*_*j*_, until only *m* units of *e*_1_ remain. This means that the net production of each entity is given by 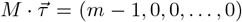, where M is the stoichiometric matrix associated to the PAC. Let us now consider a PAC witness 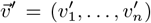, with 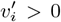 for each *i*, and scaled such that 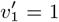 (this is always possible since scaling preserves the PAC witness inequalities). We will show that the existence of 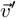 implies *m* > 1. Notice that 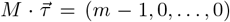. Our goal is to show that for all 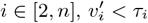

We use the graph 𝒢 defined in the previous paragraph, and more precisely the fact that the case *p* = 1 corresponds to acyclicity of 𝒢^′^ = 𝒢 \ {*e*_1_}. Without loss of generality, we can assume that a topological order of 𝒢^′^ is given by *e*_2_, *e*_3_, …, *e*_*n*_, that is to say there is no edge *e*_*i*_ →*e*_*j*_ with *j* < *i*. In particular, *e*_2_ has no incoming edge in 𝒢^′^. We compare the inflow and outflow of *e*_2_ according to flow vectors 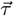 and 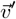.

- The inflow is the same in both cases, as 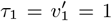 and no other incoming edge exists,
- the outflows are *σ*_2,2_τ_2_ and 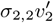 respectively,
- by construction, the net production rate with respect to 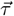 is *σ*_1,2_τ_1_ −*σ*_2,2_τ_2_ = 0, while for 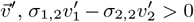 as 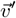 is a PAC witness.

Therefore, we have 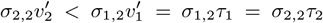, so we can conclude 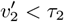. We can continue by induction with *e*_3_, *e*_4_, … : at step *i* for entity *e*_*i*_, the inflow is smaller or equal with flows from 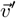 than from 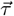 (using induction hypothesis 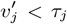 for *j* < *i*), but the net production is positive for 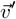 while it is 0 for 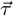. This allows us to conclude that for all *i* ∈ [2, *n*], 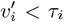. Let us call *m*^′^ the inflow of *e*_1_ according to 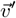, and recall that the corresponding inflow with respect to 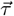 is *m*. Since every reaction producing *e*_1_ is strictly smaller in 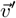 than in 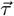, we have *m*^′^ < *m*. However, to ensure net production of *e*_1_ according to 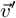, and since the outflow is *υ*_1_ = 1, we must have 1 < *m*^′^. We can conclude *m* > 1 as desired.

Now if more than one equation remains, i.e. *p* > 1 (item 2.2 of the Lemma) and *m*_*i,i*_ ≠ 1, it means that one found a subset of reactions of the cycle allowing for the net production (or consumption) of entity *e*_*i*_, in contradiction with the assumption that the cycle is minimal. An illustration of this phenomenon is given in Example 5. The existence of some *j* ≠ *i* such that *m*_*i,j*_ > 0 follows from the condition that at least one other equation than the *i*^t*h*^ remains after the substitution algorithm ends. Indeed, if we could reach an equation where *x*_*i*_ is the only remaining variable, without having substituted all the other ones, it would mean that the PAC is not strongly connected, contradicting its minimality.

###### Example 5.

*We illustrate a case with m*_*i,i*_ ≠ 1 *while two equations remain in Figure 8*.*A*. *The autocatalytic motif is not minimal, even though all entities appear only once as reactant, because it contains the core e*_2_ → *e*_3_ → *e*_1_ → *e*_1_ + *e*_2_ *(depicted in black arrows). Notice that the forked reaction e*_2_ → *e*_3_ + *e*_4_ *is not entirely in full line, because for this embedded PAC, e*_4_ *corresponds to a waste. The equations corresponding to the full system are:*

**Figure 8.**
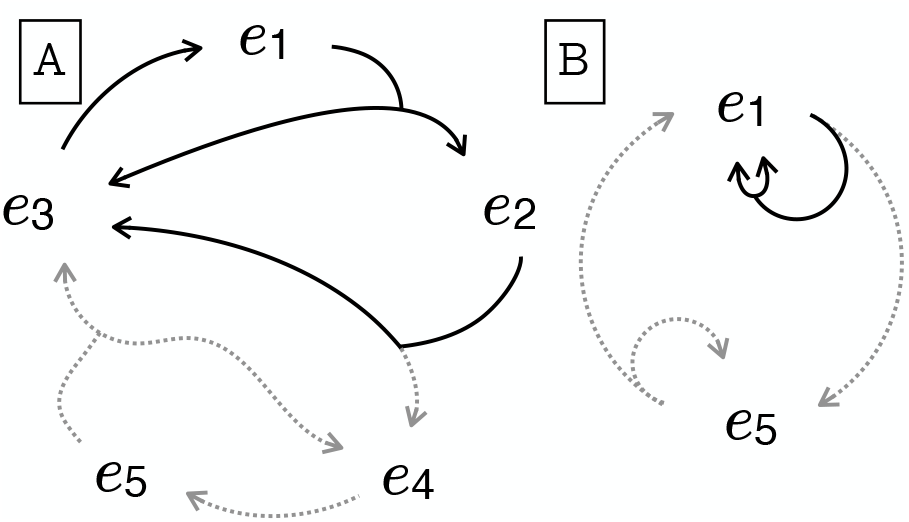
Non-minimal autocatalytic motif.

**Figure 9.**
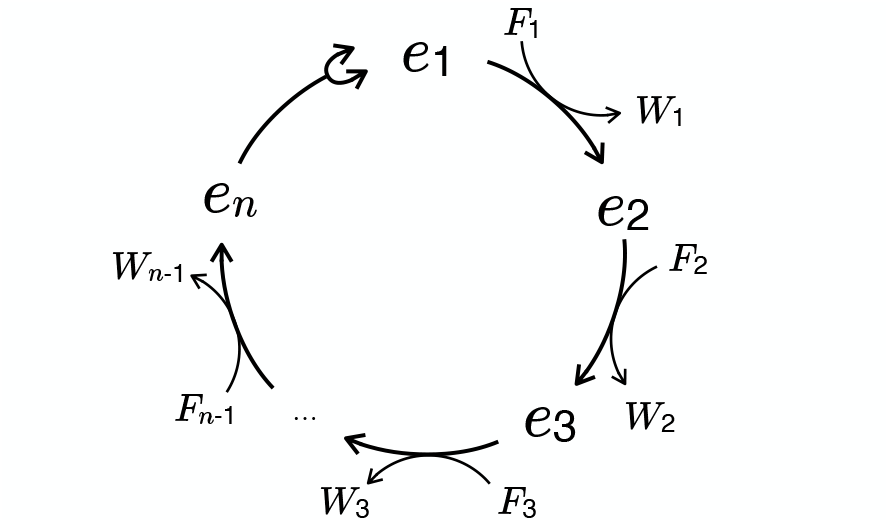
Illustration of a PAC of size n of type I, using the typology of [15].

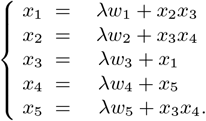

*One can substitute x*_3_, *then x*_4_ *and x*_2_ *to arrive to a system of two equations on x*_1_ *and x*_5_:

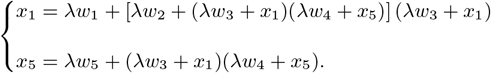

*Taking* λ = 0, *one obtains* 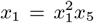 *and x*_5_ = *x*_1_*x*_5_, *thus we have m*_1,1_ = 2 *and m*_5,5_ = 1. *The appearance of x*^2^ *in the first equation betrays the existence of an inner PAC within the motif, where entity e*_5_ *is ignored. A visual interpretation is given in Figure 8*.*B, which shows the reduced cycle with the two entities e*_1_ *and e*_5_ *obtained at the end of the sequence of substitutions. Notice that entity e*_5_, *in order to self-amplify, must produce e*_1_. *However, there exists a reaction path not involving entity e*_5_ *allowing for the production of* 2 *units of e*_1_ *starting with* 1 *unit of it (full black arrows in panel B), which is why m*_1,1_ = 2. *This indeed corresponds to the PAC shown in full black arrows in panel A*.

###### Solving the reduced system

Turning back to the general proof of Theorem 2, it is enough to solve the remaining system of equations, which is of the form *x*_*i*_ = *g*_*i*_(λ, *x*_1_, …, *x*_*p*_) for all *i* ∈ [1, *p*]. We can then recover the missing *x*_*j*_ (for *j* ∈ [*p* + 1, *n*]) using the previous substitutions *x*_*j*_ = *A*_*j*_.

**Case** *p* = 1. Let us first treat the easier case where *p* = 1, i.e. we have a single variable *x*_1_ and a single equation *x*_1_ = *g*_1_(λ, *x*_1_).

###### Lemma 3.

*There exists ε* > 0 *such that for all* λ ∈ (0, *ε*), *there exists x*_1_ ∈ (0, 1) *verifying x*_1_ = *g*_1_(λ, *x*_1_).

*Proof*. Let *h*_1_(λ, *x*_1_) = *g*_1_(λ, *x*_1_) − *x*_1_. We will use the fact that *h*_1_ is continuous, and evaluate *h*_1_ at *x*_1_ = 0 and *x*_1_ = *a* for some *a* ∈ (0, 1) and for a suitable λ. We will conclude the proof using the intermediate value theorem.

First of all for any λ > 0 and *x*_1_ = 0, we have *h*_1_(λ, 0) > 0, since by construction *g*_1_ is built from positive terms using functions preserving positivity, and furthermore contains a term of the form λw_1_ > 0 not depending on *x*_1_.

Now, let us consider the case *x*_1_ = *a* for any *a* < 1. We have *h*_1_(λ, *a*) = *g*_1_(λ, *a*) −*a*.

By Lemma 2, for λ = 0 we have *g*_1_(0, *a*) = *a*^*m*^ with *m* > 1. Since *a* < 1 and *m* > 1, we have *a*^*m*^ < *a*. Thus, *h*_1_(0, *a*) < 0. Since the function λ → *h*_1_(λ, *a*) is continuous, there exists *ε* > 0 such that for all λ ∈ (0, *ε*), we have *h*_1_(λ, *a*) < 0. We can conclude (intermediate value theorem with respect to *x*_1_) that for all λ ∈ (0, *ε*), there exists *x*_1_ ∈ (0, *a*) such that *h*_1_(λ, *x*_1_) = 0.

Therefore, choosing any λ < *ε* and applying this lemma gives a solution to the system.

**Case** *p* > 1. If *p* > 1, we will solve the system by constructing a sequence of partial continuous functions *f*_*i*_ : ℝ^*i*^ → ℝ (1 ≤ *i* ≤ *p*) and *ε* > 0 such that:

- if *x*_1_, …, *x*_*i*−1_ ∈ (0, 1) (resp. [0, 1)) and λ ∈ (0, *ε*) (resp. [0, *ε*)), then *f*_*i*_(λ, *x*_1_, …, *x*_*i*−1_) is defined, and its image lies in (0, 1) (resp. [0, 1)),
- if there exists λ, *x*_1_, …, *x*_*p*_ such that for all *i, x*_*i*_ = *f*_*i*_(λ, *x*_1_, …, *x*_*i*−1_), then we have a solution of the wanted equations.

We define *f*_*i*_ by induction, starting with *f*_*p*_.

Let us consider the equation *x*_*p*_ = *g*_*p*_(λ, *x*_1_, …, *x*_*p*_). Let *h*_*p*_(λ, *x*_1_, …, *x*_*p*_) = *g*_*p*_(λ, *x*_1_, …, *x*_*p*_) − *x*_*p*_.

Our goal is to define *f*_*p*_ as a value of *x*_*p*_ yielding *h*_*p*_ = 0. Thus, we have to show that such a root of *h*_*p*_ exists.

###### Lemma 4.

*For any x*_1_, …, *x*_*p*−1_ ∈ (0, 1), *there exists ε*_*p*_ *such that for all* λ ∈ (0, *ε*_*p*_), *there exists x*_*p*_ ∈ (0, 1) *verifying h*_*p*_(λ, *x*_1_, …, *x*_*p*−1_, *x*_*p*_) = 0.

*Proof*. As before, we will use the fact that *h*_*p*_ is continuous (by construction of *g*_*p*_), and look at the values of *h*_*p*_ for *x*_*p*_ = 0 and *x*_*p*_ = *a* < 1.

First of all for *x*_*p*_ = 0, for any λ, *x*_1_, …, *x*_*p*−1_ > 0, we have *h*_*p*_(λ, *x*_1_, …, *x*_*p*−1_, 0) > 0, by construction of *g*_*p*_.

Now, let us consider the case *x*_*p*_ = *a* for some arbitrary *a* <1. We have *h*_*p*_(λ, *x*_1_, …, *x*_*p*−1_, *a*) = *g*_*p*_(λ, *x*_1_,. .TI., *x*_*p*−1_, *a*) − *a*. For λ = 0, we have 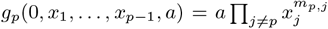 by Lemma 2.

By Lemma 2, we have in addition *m*_*p,j*_ > 0 for some *j* ≠ *p*. Thus, for *x*_1_, …, *x*_*p*−1_ < 1, we have 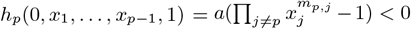. Since *h*_*p*_ is continuous, there exists *ε*_*p*_ such that for all λ ∈ (0, *ε*_*p*_), we have *h*_*p*_(λ, *x*_1_, …, *x*_*p*−1_, *a*) < 0. By the intermediate value theorem, for all λ ∈ (0, *ε*_*p*_), there exists *x*_*p*_ ∈ (0, *a*) such that *h*_*p*_(λ, *x*_1_, …, *x*_*p*−1_, *x*_*p*_) = 0. Here we could actually have taken *a* = 1, but we give the scheme that will be used throughout the rest of the induction.

We can thus define *f*_*p*_(λ, *x*_1_, …, *x*_*p*−1_), as the least value of *x*_*p*_ such that *h*_*p*_(λ, *x*_1_, …, *x*_*p*−1_, *x*_*p*_) = 0, as soon as the conditions of the lemma are met. Moreover, *f*_*p*_ is continuous as well, since it is defined as the first root of a continuous function with non-zero boundary conditions.

We now get rid of equation *x*_*p*_ = *g*_*p*_ by replacing *x*_*p*_ with *f*_*p*_(λ, *x*_1_, …, *x*_*p*−1_) in all the other equations. We then have a system of the form *x*_*i*_ = *g*_*i*_(λ, *x*_1_, …, *x*_*p*−1_, *f*_*p*_(λ, *x*_1_, …, *x*_*p*−1_)) for all *i* ∈ [1, *p* − 1].

Notice that replacing *x*_*p*_ with *f*_*p*_ allows us to carry out the previous construction for *f*_*p*−1_, as the only property asked of the various *x*_*j*_ other than the current *x*_*p*−1_ under consideration is to be in (0, 1), and that is the case for *f*_*p*_. Notice that *x*_*p*_ is now a function of λ, *x*_1_, …, *x*_*p*−1_, but this is not a problem, as long as it remains in (0, 1) (allowing for extremal cases if some arguments are extremal as well), the construction can be carried on.

We can thus iterate this construction, and obtain a sequence of functions *f*_*i*_ as wanted. At each step, the *ε*_*i*_ can be chosen to be the minimum of the *ε*_*i*+1_ obtained at the previous step and the *ε* needed at the current step. This guarantees that all *f*_*j*_ for *j* ≥ *i* are defined for all λ < *ε*_*i*_.

Continuing this construction, we will end up with a sequence of functions *f*_*i*_ such that for all λ < *ε*, there exists *x*_1_, …, *x*_*p*_ such that *x*_*i*_ = *f*_*i*_(λ, *x*_1_, …, *x*_*i*−1_) for all *i* ∈ [1, *p*]. This will give us a solution to the system of equations, and thus a solution to the original problem. Indeed, we only need to choose some λ < *ε*_1_, *x*_1_ = *f*_1_(λ), *x*_2_ = *f*_2_(λ, *x*_1_), etc. to obtain a solution to the system.

As described earlier, we can infer the values of *x*_*p*+1_, …, *x*_*n*_ from *x*_1_, … *x*_*p*_ by using the previous substitutions *x*_*j*_ = *A*_*j*_. From this, we can finally compute the concentration vector 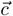 such that 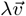 is the flow vector induced by 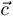.

As explained in Corollary 1, this is sufficient to show that any PAC is CAC, as we can start with a PAC witness 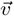 and apply this construction to find a CAC witness 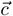.

#### 4. On the degree of feasibility of PACs

In this section we investigate how thermodynamic constraints affect the degree to which a PAC can be a CAC, in the sense of exploring quantitatively and analytically how such constraints, though not being able to prevent the feasibility of PACs (see Corollary 1), nonetheless affect the limitations toward instantiating a PAC in the concentration space. We will focus on a specific topology of type I PACs, following the terminology of [15]. We will consider a PAC of size *n* with a single fork, while reactions other than the fork may consume food and produce waste. These assumptions are for illustration purposes, and we believe the phenomena that we will describe here occur more generally. Thus, the PAC considered is of the following form, where *e*_*n*_ → *e*_1_ + *e*_1_ is the forked reaction:

Following the notations of the proof explained in Appendix 3, one would end up considering the following system of equations:

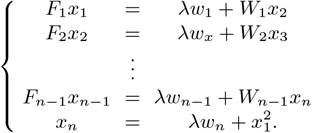

*F*_*i*_ (respectively *W*_*i*_) is the exponential of the chemical potential of food (respectively waste) *i*. The substitution algorithm from Section 3 allows us to inject the (*i* + 1)^th^ line into the *i*^th^ until one finally obtains a single equation in *x*_1_:

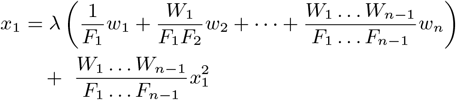

rewritten as:

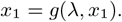

For this particular PAC example, the dependence of λ with respect to other parameters can be made explicit:

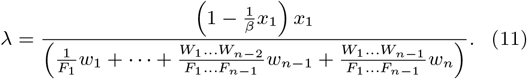

It should be noted that the condition λ > 0 imposes *x*_1_∈ (0, *β*) with 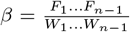. Notice that if all food and waste variables are set to 1, one recovers the interval of the general proof of Appendix 3 which was (0, 1). *R*egardless, the PAC can be realized for any value of food and waste potentials.

Equation (11) shows that the solution λ has non-trivial dependencies on food, waste and PAC species chemical potentials, activation barriers of reactions and the witness 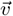. To facilitate the discussion, let us assume that the witness 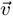 is the one allowing for a unitary production of all PAC species (such a witness always exists because the stoichiometry matrix of a PAC is square and invertible, see discussion in [15, SI, proposition 2]). Hence, because the witness 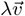 can be instantiated in the concentration space, λ can be interpreted as the amplitude of the PAC, i.e. the level up to which it can effectively produce its entities, or its degree of feasibility, relative to the case of a unitary production rate of PAC entities.

Notice that both the numerator and the denominator of Equation (11) depend on terms of the form Π_*i*_ *W*_*i*_/ Π_*j*_ *F*_*j*_. The denominator becomes smaller as the food variables grow larger relative to the waste variables. As far as the numerator is concerned, we plot in Figure 10 the function θ_*β*_(*x*_1_) = (1 − *x*_1_/*β*)*x*_1_ for two values of *β* (namely *β* = 1 and *β* = 2). θ_*β*_ is an inverted parabola which is positive for *x*_1_ ∈ [0, *β*] and reaches it maximum at *x*_1_ = *β*/2, the maximum being *β*/4. Hence, the numerator of Equation (11) (taken as a function of *x*_1_) can get larger as *β* gets larger, i.e. as food variables get large in comparison to waste variables. We thus conclude that λ can reach higher values when the food variables are globally large in comparison to waste variables, which from now on is referred to as a favorable environment.

**Figure 10.**
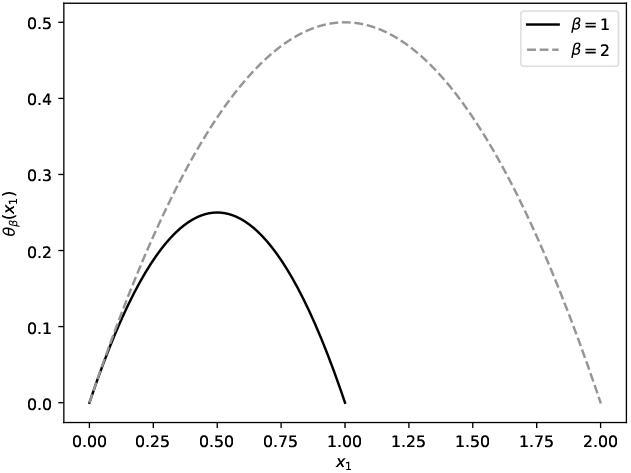
The numerator of Equation (11), θ_*β*_(*x*_1_) = (1 − *x*_1_/*β*)*x*_1_ as a function of *x*_1_ for *β* = 1 and *β* = 2.

A favorable environment can always be reached by choosing appropriate food and waste concentrations. However, it should be recalled that the variables appearing in Equation (11) are related to the free energy and concentration of entities through the relation 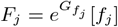 (illustrated for the *j*^th^ food species).

We have seen that the PAC realization is easier when *F*_*j*_ ‘s are large relative to W_*i*_ ‘s. *R*ecall that 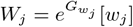. If one waste free energy is increased by *δG*, then this corresponds to a multiplication of the corresponding *W*_*j*_ by *e*^*δG*^, which will have to be compensated for by similar variations in the concentrations of food and/or waste entities. Thus, slight changes in the free energy values are exponentially passed on the concentration space. This highlights how PACs whose food and waste free energies are unfavorably distributed can be hard to instantiate in the concentration space in practice.

Similar conclusions can be reached regarding the concentrations of PAC entities. Notice that in Equation (11) λ depends only on *x*_1_ in the numerator shown in Figure 10. This shows that *x*_1_ is upper bounded, so the concentration space of entity *e*_1_ associated to λ > 0 (i.e. the PAC running) increases its size as its free energy decreases. Notice once again that a favorable environment (i.e. large values of *β*) is associated with a wider interval of *x*_1_ allowing λ > 0.

We conclude that regions of the concentration space where the PAC efficiently runs (i.e. runs with large values of λ) always exist if all concentrations can be chosen freely, but that free energies unfavorably distributed among food, waste and PAC entities will significantly restrict their volume.

On similar grounds, one can study the impact of activation barriers on the feasibility of PACs. Suppose that all reactions of the PAC get penalized via an increase of their activation barriers, such that 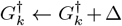 with Δ > 0 for all *k* ∈ [1, *n*]. This will in turn multiply the denominator of Equation (11) by a global factor *e*^Δ^, and, as a consequence, multiply λ by *e*^−Δ^ < 1. This can be compensated by choosing a more favorable environment, i.e. by increasing food concentrations and diminishing waste concentrations, such that *β* increases. Thus, the regions of the concentration space associated with large values of λ are tightened.

1 In this formulation, we set *RT* = 1, which can be done without loss of generality, as this is simply equivalent to expressing Gibbs free energies in *RT* units. In the following, the notation *R* will stand for a reaction, and the gas constant will not be referred to again. Similarly, other physical constants will be ignored in order to set a dimensionless framework for simplicity. Such quantities could be reintroduced without affecting the results presented in this work.

2 This is actually an instance of 2SAT which is a simpler problem, but it is just used here to illustrate the construction

